# Genomes2Genes: A Progressive Cactus Gene Analysis Pipeline

**DOI:** 10.64898/2026.11.08.765189

**Authors:** Sierra C. Sanders, Claudio Casola

## Abstract

Whole-genome alignments are fundamental resources in comparative genomics, because they place genes and their surrounding sequences in a shared evolutionary context. Progressive Cactus aligns large numbers of genomes without a reference genome and reconstructs ancestral sequences along the input phylogeny. However, its Hierarchical Alignment Format output has mainly been used for visualization, phylogenomics, and pangenome studies, and gene-level analyses of these alignments remain difficult. Here, we present Genomes2Genes, a pipeline that converts Progressive Cactus graph-based alignments into validated coding sequence alignments of extant species and their reconstructed ancestors. The pipeline extracts exon alignment blocks, handles atypical genomic jumps, stitches exons into transcripts, and reports start codons, stop codons, and frameshifting indels across species, with validation steps throughout and a base-by-base audit of the final alignments. It catches both biological and technical artifacts and can be run on any part of the genome. We anticipate that this pipeline will be useful in identifying and removing problematic regions used in the potential reconstruction of species or gene phylogenies. We developed and tested the pipeline on a whole-genome alignment of 21 Brassicaceae species and 2 outgroups and found that, among 45,491 lineage-specific genes, the pipeline supported 22,376 candidate *de novo* genes, of which 1,811 had complete evidence of enabler mutations. Genomes2Genes is publicly available and extends whole-genome alignments to gene-level questions in comparative genomics and evolutionary biology.

## Introduction

Genome sequence alignments are critical resources to identify and analyze genetic variation across populations and species, with a growing impact in evolutionary biology, medical genetics, and agriculture (Sullivan et al. 2023; Wu et al. 2023; Zoonomia Consortium 2020). The rapidly increasing number of high-quality genomes (Lewin et al. 2018; Rhie et al. 2021) has introduced a strong demand for bioinformatic tools to produce large-scale whole-genome alignments (WGAs), including pangenomes (Hickey et al. 2024; Liao et al. 2023), both effectively and with limited computational time. Progressive Cactus (PC) (Armstrong et al. 2020) has become one of the most widely used tools to generate WGAs. A major advantage of this program is that it builds reference-free whole-genome alignments, which therefore are not restricted to the sequences of a single reference genome (Armstrong et al. 2020), overcoming one of the most crucial biases of other gene and genome alignment programs. Additionally, the runtime of PC scales linearly with the number of genomes and their sizes (Armstrong et al. 2020).

Developed to address the challenges of aligning hundreds of large genomes, including those of the Zoonomia and Bird 10,000 Genomes projects (Andrews et al. 2023; Armstrong et al. 2020; Christmas et al. 2023; Feng et al. 2020; Sullivan et al. 2023; Zoonomia Consortium 2020), PC has proven successful in studies of human genetics (Sullivan et al. 2023), selective constraint in birds (Feng et al. 2020) and placental mammals (Christmas et al. 2023), the genomic architecture of teleost fish (Song et al. 2026), *de novo* gene origination in *Drosophila* (Peng and Zhao 2024), and the first comprehensive analysis of aligned telomere-to-telomere ape pangenomes (Yoo et al. 2025). PC has also been used to analyze plant genomes, which are notoriously difficult to effectively align at scale because of their rapid sequence divergence and erosion of synteny across species (Song et al. 2024; Wendel et al. 2016). Studies in potato have demonstrated its capacity to handle the unique genomic attributes found across plants (Tang et al. 2022; Wu et al. 2023; Zhang et al. 2025). One of the most important yet underutilized features of PC is its reconstruction of ancestral sequences during multiple genome alignment, which offers a unique and critical avenue for investigating questions about genome evolution in a phylogenetic context, because the ancestral state of a sequence can be compared directly with its extant state (Armstrong et al. 2020).

While PC addresses many of the deficits of other WGA tools, several weaknesses have hampered its wider applications, in particular in the analysis of non-genic regions. One of these, which is shared across similar tools, is its dependence on an input guide tree, which can affect the resulting alignment when the species phylogeny is uncertain. This was determined to not present as a significant issue in minor changes to the trees through a four alignment test study from 48 avian species with different guide trees, with 98.5% of aligned pairs identical on average (Armstrong et al. 2020). However, the dependence on the quality of the guide tree cannot be completely eliminated and has been shown to be robust only to small changes in the input trees, as described in the creation of Minigraph-Cactus, analyzing pangenomes (Hickey et al. 2024).

PC alignments also vary across the genome, because alignments of the same species agree more strongly in coding exons than across the genome as a whole (Armstrong et al. 2020), likely because coding regions are evolutionarily conserved compared to intronic and intergenic sequences.

Despite these weaknesses, PC is one of the most accurate tools currently available (Armstrong et al. 2020; Earl et al. 2014; Paten et al. 2011) and offers many avenues for examining the different aspects of a genome. Its primary output is a Hierarchical Alignment Format (HAL) file, which is a graph-based representation of the multiple genome alignment (Hickey et al. 2013). Resources such as the UCSC Genome Browser (Casper et al. 2026; Hickey et al. 2013) and HAL tools (Hickey et al. 2013) adapt the HAL file so that it can be visualized, but their applications are limited. Beyond phylogenomic or pangenome studies (Hickey et al. 2024; Yoo et al. 2025), HAL-based tools address specific tasks, including the construction of synteny blocks with halSynteny (Krasheninnikova et al. 2020), the projection of gene annotations with the Comparative Annotation Toolkit (CAT) (Fiddes et al. 2018), and the mapping of regulatory element orthologs with HALPER (Zhang et al. 2020). These also present new challenges by limiting the output from these tasks to only a pairwise comparison per run (Hickey et al. 2013; Krasheninnikova et al. 2020; Zhang et al. 2020) and CAT re-introduces reference bias (Fiddes et al. 2018). Efficient ways to obtain validated gene-level alignments of extant and ancestral sequences from these files, without removing any of the unique advantages from using PC, remain scarce. The HAL format and toolkit were initially designed to address issues that arose with reference-free whole-genome alignments, particularly their indexing (Hickey et al. 2013). The graph-based structure of the HAL files fixes these issues and was also created to compress the output alignments in anticipation of large-scale genome alignment analyses. The limited command line utilities from HAL tools also deal with indexing this file format and extracting comprehensive information across many species (Hickey et al. 2013).

Here, we present the Genomes2Genes pipeline to address this gap. It works in conjunction with the HAL output file and other established PC toolkits to convert and analyze the alignments in ways that allow the investigation of many research questions, rather than being limited to phylogenomics or building pangenomes. Our custom scripts build on HAL tools and mafTools (Earl et al. 2014) and include 17 steps that filter and break down the different elements of these alignments, thereby extending the range of applications of one of the most important bioinformatic tools in comparative genomics. We constructed the pipeline using 21 species from the plant family Brassicaceae and 2 outgroups, which permitted robust testing of each component during its development. The study focused on developing a computational genomics pipeline to accurately and efficiently identify *de novo* genes in this family. *De novo* genes arise from non-coding sequences in the genome that gain the ability to be transcribed and translated through enabler mutations (Van Oss et al. 2019; Zhang et al 2019; Zhao et al. 2014). The mutations can create an open reading frame (ORF), create start codons, and remove premature stop codons (Zhang et al. 2019). These genes tend to be short in length and have high turnover rates, making it extremely difficult to identify them (Casola et al 2025). The most concrete evidence for confirming *de novo* origin is using a combination of homology and synteny across species (Vakirlis et al. 2020; Zhang et al. 2019), the latter requiring the PC alignment processing and validation implemented in Genomes2Genes pipeline. The presence of ancestral sequences is also a key factor in pinpointing the enabler mutations and origin of these genes (Vakirlis et al. 2020). Working with plant genomes, which present well-known hurdles including polyploidy, highly repetitive sequence regions, whole-genome duplications, and fast-eroding synteny (Claros et al. 2012; Walden et al. 2020; Wendel et al. 2016), allowed us to address these challenges as the pipeline was being assembled. In addition to evaluating the rigor of the PC alignments themselves, the pipeline covers the entire filtering and evaluation processes, from a HAL output file to the extraction of Multiple Alignment Format (MAF) blocks and the analysis of individual base-pair alignments between species and genes.

### Genomes2Genes Overview

The pipeline is organized around four sections: Extraction, Filtering & Conversion, Assembly, and Analysis. The first section of the pipeline, Extraction, uses the established HAL and MAF comparative genomic toolkits (Earl et al. 2014; Hickey et al. 2013). The HAL file is converted into a MAF file with HAL tools, which requires a reference species to be selected. This does not remove the reference-free aspect of the original alignment, because the reference is used solely to extract portions of the alignment. To investigate a cross-species research question, multiple MAF files need to be created with each relevant species selected as the reference. A Browser Extensible Data (BED) file containing the genomic coordinates of the sequence regions of interest is used with the mafExtractor program from mafTools to isolate and extract these local alignment blocks, and every exon extracted is output in its own file. Any extraction that fails or returns no sequence stops the process to prevent downstream errors.

To ensure structural and genomic integrity of these extractions, the second section of the pipeline, Filtering & Conversion, identifies the species present in each extracted MAF block per file and indicates whether any base pairs are missing between blocks due to low alignment quality and, if so, from which species and where in the alignment (**Fig. 1**). It then handles ‘atypical genomic jumps’, namely regions of a species or ancestral sequence whose consecutive blocks lie on different chromosomes or strands, break collinearity with the reference genome, or lie more than 50 kb apart on the same chromosome. The computational flaws leading to these jumps are due mostly to the coordinate-based system used by mafTools and mafExtractor, which can pull non-contiguous sequences because of the graph-based nature of the original HAL alignment. However, chromosomal jumps could also be due to errors in the annotation and sequence assembly of the genomes. By default, the MAF blocks of each species within an exon are grouped by chromosome and strand, so the Genomes2Genes pipeline identifies the species that fall into any of those categories and removes them from the entire exon for each file (**Fig. 1**).

**Figure 1.**
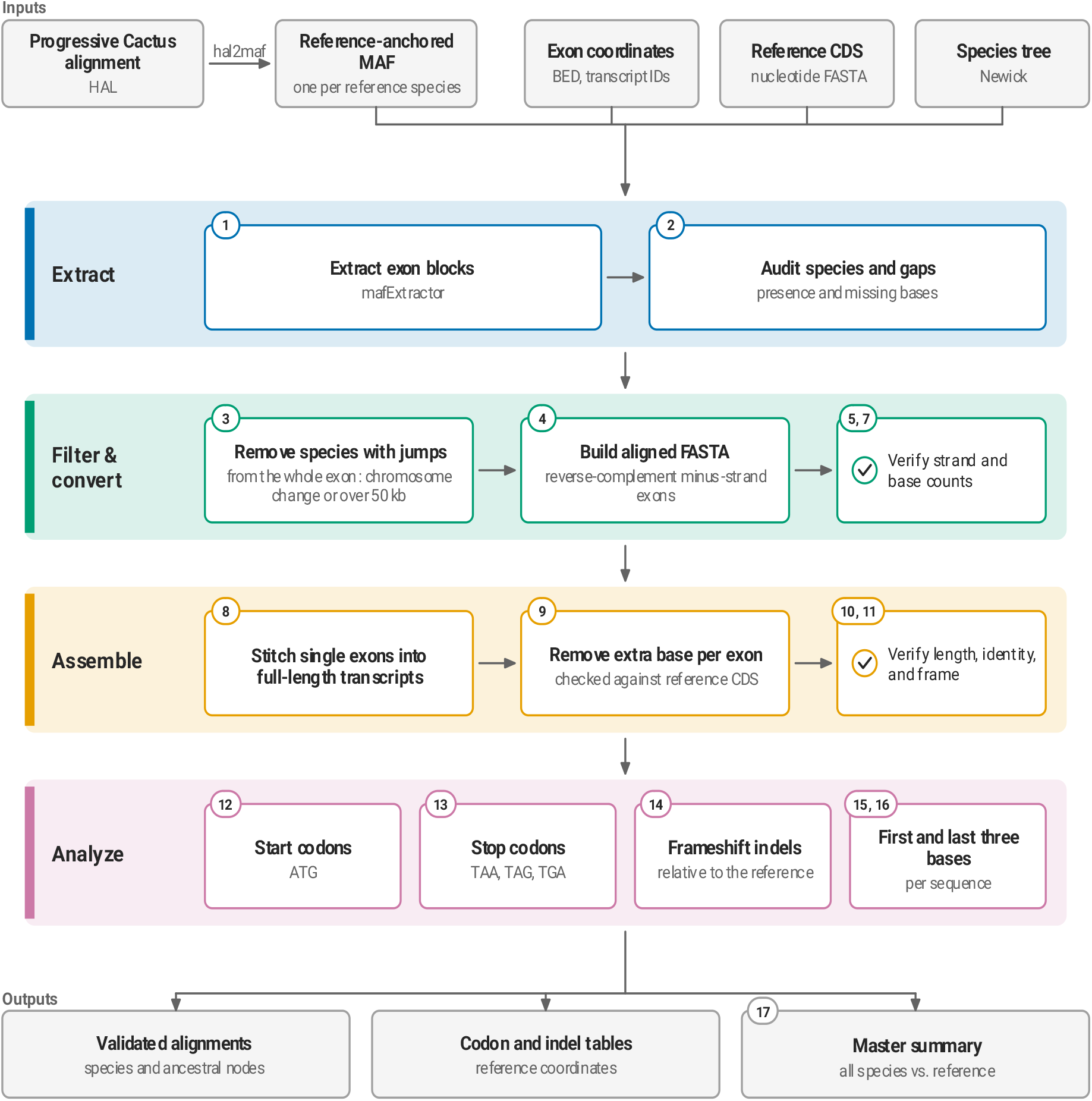
Workflow of Genomes2Genes. A Progressive Cactus whole-genome alignment is exported as a reference-anchored MAF and processed in four sections: Extraction, Filtering & Conversion, Assembly, and Analysis. The result is validated codon-level alignments of extant species and reconstructed ancestors, additionally including summaries of start codons, stop codons, and frameshifting indels. Numbers mark pipeline steps; check marks indicate validation steps. Each section is described in the Genomes2Genes Overview.

Each exon is then converted into an aligned FASTA file by concatenating the MAF blocks of each species, with gaps added for any species absent or removed from a block so that all sequences remain the same length (**Fig. 1**). Bases that a species skips between consecutive blocks are unaligned sequence that did not score high enough on PC’s alignment criteria and are deliberately not added back in, because filling them would force gap columns into the reference sequence and shift the frame of the other species relative to it. The sequences are also reverse-complemented when the reference exon is on the negative strand based on the BED file information, because mafTools always assumes that all coordinates entered for extraction are on the positive strand. Strand orientation is then verified by local alignment against the original MAF sequences, and nucleotide counts are compared across the raw, filtered, and converted files.

The third section, Assembly, then stitches the exons of each transcript together, in transcript order, into a single alignment using the transcript IDs of the reference species provided in the BED file (**Fig. 1**). Multi-exon sequences must be combined to produce the full-length transcript because each exon is written to a separate output file by mafExtractor.

Because mafExtractor treats the end coordinate of each exon as inclusive, every extracted exon contains one extra base beyond the annotated exon boundary. These are removed at their known reference positions from the BED coordinates (the last base of each exon on positive-strand genes and the first base on negative-strand genes), rather than by matching sequence, so that every species is cut at the same position even when the extra base is identical to its neighbor.

Columns in which the reference has a gap are also removed, projecting every species onto the reference coding sequence (CDS). A trimmed alignment is written only if its reference sequence matches the reference CDS in that species’ nucleotide FASTA file; every other alignment is excluded and listed in a report with the reason. The final alignments are then validated against the reference in length and identity using the BED file, and the reading-frame periodicity of each transcript is checked to ensure it has been maintained.

With the sequences for the gene region of interest filtered and validated, the alignment can be analyzed across species and ancestral sequences to answer evolutionary and biological questions. Because the pipeline was originally developed to identify mutations that enable the formation of longer open reading frames (ORFs) in *de novo* gene birth, known as enabler mutations (Knowles and McLysaght 2009; Van Oss and Carvunis 2019), the final section of the pipeline, Analysis, includes scripts that locate start codons (ATG) and stop codons (TAG, TAA, and TGA), reading each species in its own frame from its first aligned base and reporting them with respect to the reference coordinates for comparability (**Fig. 1**). It also identifies biological indels that cause frameshifts, defined as runs of gaps whose length is not a multiple of three, which can represent enabler mutations. For species reduced to a single row during the exon stitching process, indels that arise only from the exons of their dropped rows are removed, but every removed indel is still recorded. Finally, the pipeline extracts the first three and last three base pairs of each sequence in the alignment, which allows for comparison of the start and end of each sequence across species.

All steps of the pipeline generate a summary CSV or TSV file compiling the information gathered at that step, the validation steps report their results in the run log, and after the final step, a master summary file combines the results for every species relative to the reference. A single script runs all steps in order for one reference species at a time. Before starting, it checks that all inputs and required tools are available, and it records the status of each run (running, succeeded, or failed at a specific step) together with its provenance. Runs can be resumed from any step, and because every output read by a later step is written in a fixed order, re-running the pipeline on the same inputs produces the same files. An end-to-end audit then checks the final alignments base by base, confirming that every expected base is present at its column, that no unexplained bases are present, and that the reference sequence equals its CDS. All relevant scripts and files are available at https://github.com/sierras64/Genomes2Genes-Brassicaceae.

Genomes2Genes requires HAL tools and mafTools used from Progressive Cactus v3.2.1, GNU parallel 20260722 (Tange 2011), and Python v3.10.20 with Biopython v1.87 (Cock et al. 2009) and pandas v2.3.3 (McKinney 2010). A run takes a working directory, the reference species, a BED file of exon coordinates with transcript IDs, the reference CDS FASTA file, the MAF file, and a Newick tree. Each step can be run on its own, and each is documented in the repository.

### Initial Test in Brassicaceae

The 21 Brassicaceae species and 2 outgroups (*Tarenaya hassleriana* and *Theobroma cacao*) used to create and test the pipeline included the model plant *Arabidopsis thaliana* as well as important crop species such as *Brassica rapa*. The whole-genome alignment of these species, produced with PC v3.2.1, is, to our knowledge, one of the largest whole-genome alignments of species across a plant family. Applying PC to a clade of species with rapidly evolving genomes also allowed us to test possible limitations of PC. Our alignment showed the expected trend of decreased coverage between increasingly distant species along the phylogeny, and an average coverage of 33% in Brassicaceae and the two outgroup species (**Fig. 2**).

**Figure 2.**
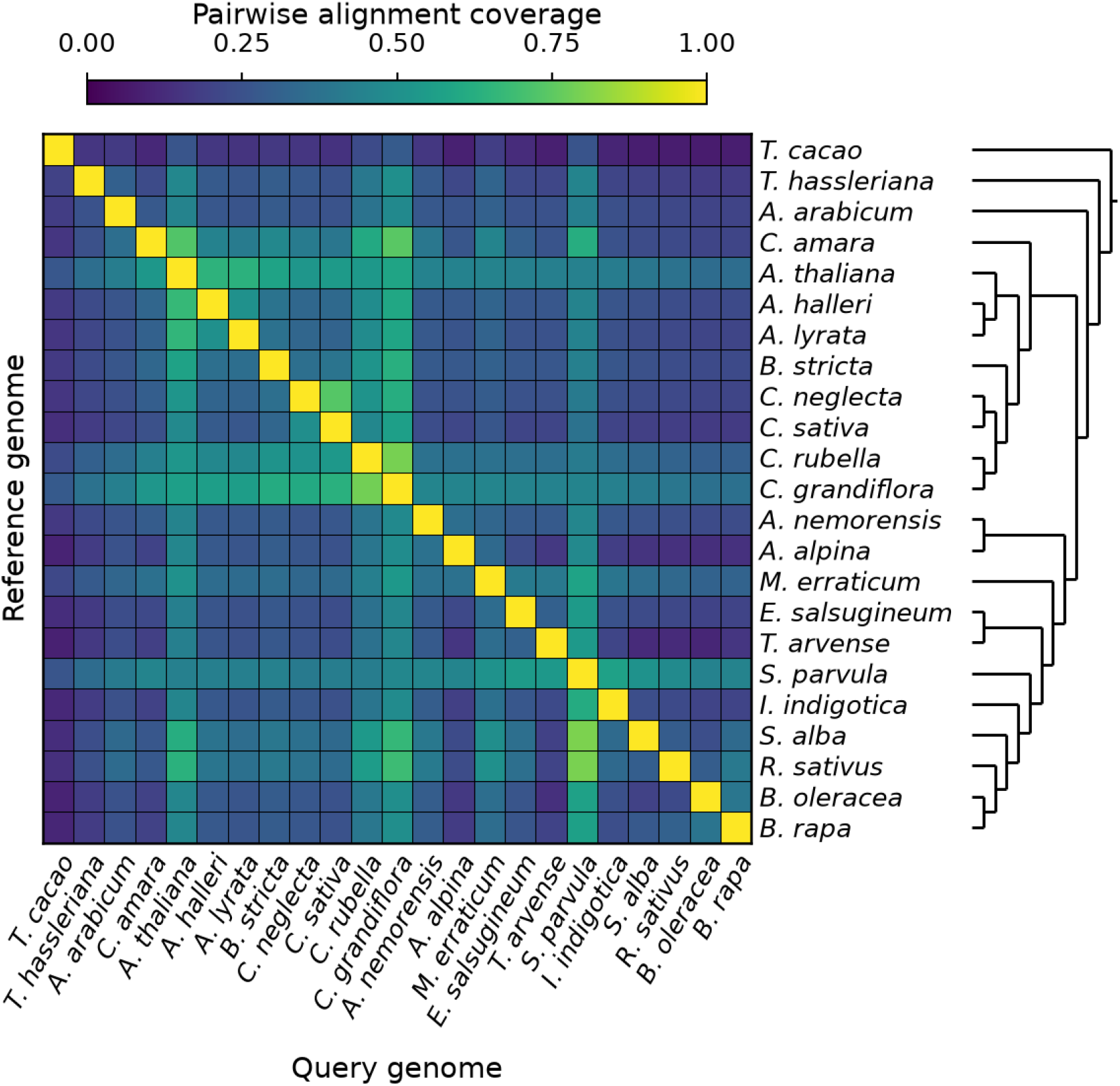
Pairwise whole-genome alignment coverage of the 23-genome Progressive Cactus alignment (21 Brassicaceae species and two outgroups *Tarenaya hassleriana* and *Theobroma cacao*). Each cell is the number of bases aligned one-to-one between the row genome and the column genome, taken from the coverage table generated by PC hal2maf and anchored on the row genome, divided by the smaller genome size of the two. Each row comes from a different reference-anchored MAF, so the matrix is asymmetric; a cell and its mirror across the diagonal one-to-one coverage line measure the same pair of species’ coverage in opposite directions. Species are ordered by the cladogram at right. The color scale is linear.

These coverage files can be generated along with the MAF file for each species. When calculating the coverage on the Brassicaceae run, both genome size and polyploidy were accounted for. A total of 45,491 lineage-specific genes were identified across these species using the LINGUA pipeline [https://github.com/CasolaLab/LINGUA], and these lineage-specific genes were run through the Genomes2Genes pipeline. A MAF file was generated with each species as the reference and was used to extract candidate genes for each species. This large set of genes allowed us to identify issues with the HAL tools and mafTools programs, as well as previously mentioned biological and genomic hurdles, which we were then able to resolve with the pipeline, additionally addressing the atypical genomic jumps within exons. All candidate genes were successfully run through the full pipeline, and start codons, stop codons, and indels were identified across the alignments for each gene. Because lineage-specific genes can also result from homology detection failure or duplication (Casola 2018; Vakirlis et al. 2020; Weisman et al. 2020), synteny is required to support a *de novo* origin. Of the 45,491 lineage-specific genes, 22,376 (49.2%) were revealed as candidate *de novo* genes, with 1,811 (8.1% of candidates) having complete evidence of enabler mutations indicating high-confidence *de novo* gene status, based on the outputs from the pipeline (**Fig. 3**). The remaining candidates lacked sufficient synteny data, a known issue in analyzing plant genomes (Song et al. 2024), to be definitively classified as high-confidence *de novo* genes and were therefore termed putative *de novo* genes.

**Figure 3.**
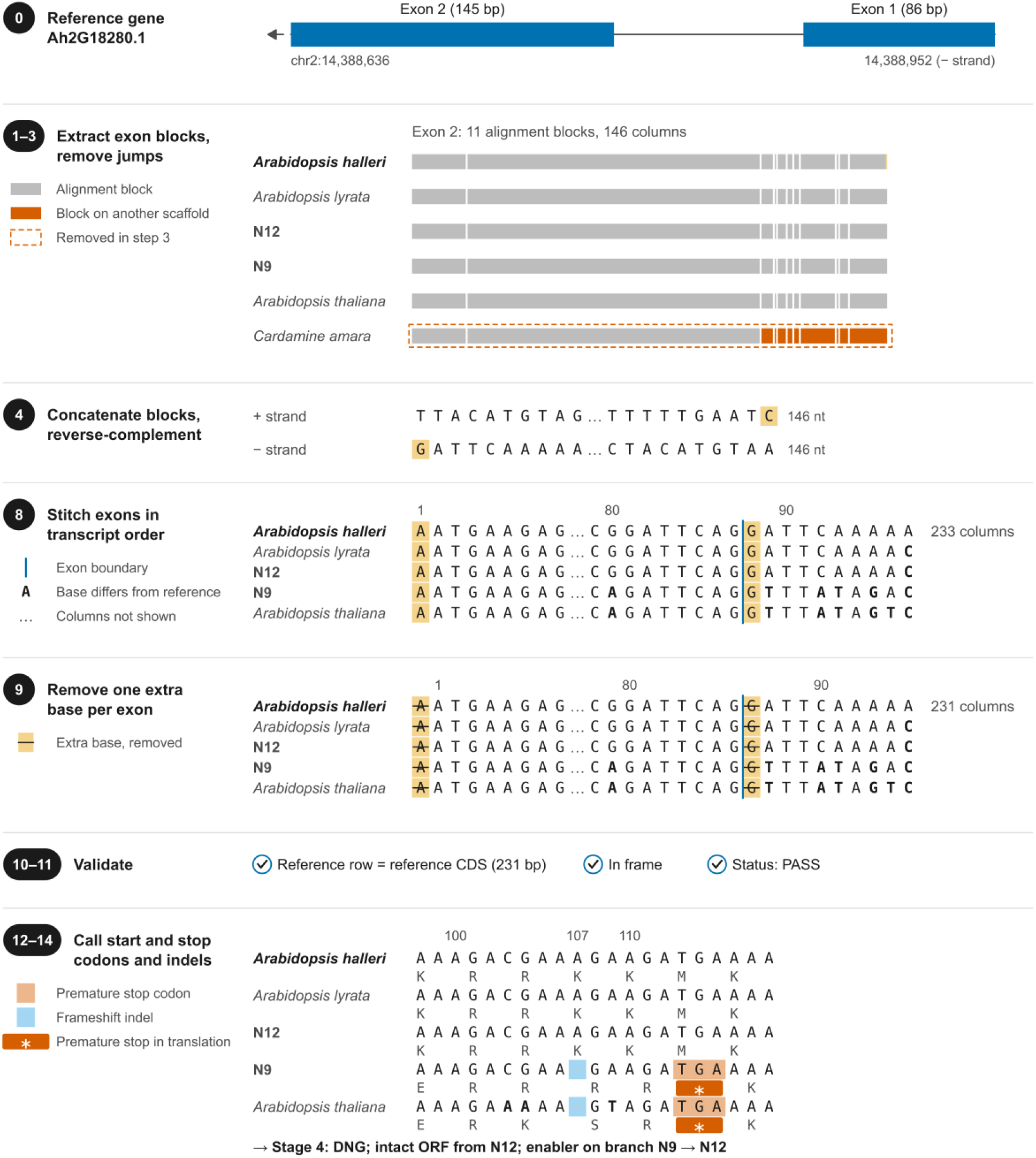
*A. halleri* gene (*Ah2G18280*.*1*) from the Brassicaceae run. Steps and symbols are indicated by the legends on the left of each segment. This gene has two exons on the minus strand of chromosome 2 (86 and 145 bp). Exon 2 is returned as 11 blocks (146 columns); *Cardamine amara* subsp. *amara* switches from scaffold *JBANAX010000077*.*1* to *JBANAX010000114*.*1* at block 3 and is removed in steps 1–3 from exon 2. Exon 1 is not shown here. N9 and *A. thaliana* carry a 1-nt deletion at position 107, after which a TGA stop codon falls in frame at positions 113–115 (columns 98–118 shown). The ORF is intact from N12, so the enabler lies on the N9→N12 branch for *A. halleri*. For further explanation of each step see the Genomes2Genes Overview.

The *QQS* gene (Li and Wurtele 2015) in *A. thaliana* is one of the best-characterized candidate *de novo* genes in plants (Arendsee et al. 2019) and has been experimentally determined to be involved in starch metabolism (Li et al. 2009). *QQS* has been classified as a *de novo* gene (Silveira et al. 2013), but its origin via this mechanism remained unverified due to the lack of synteny conservation data. Our PC alignment provides the first alignment-based syntenic evidence that it could have arisen *de novo* (**Fig. 4**). Its ORF is fully intact only in *A. thaliana*, whereas the syntenic sequence in its sister species *A. halleri* and *A. lyrata* and in two ancestral sequences contains a 100-bp gap and a premature stop codon in exon 1 (**Fig. 4**). Although the lack of synteny conservation in further species prevents the formal classification of *QQS* as a high-confidence *de novo* gene, our PC alignment data strongly support its classification as a putative *de novo* gene. In contrast, the *SWK* gene (Jin et al. 2025), previously identified as a *de novo* gene using homology searches across species of the same family, showed no conserved synteny with any species or ancestor in our alignment of the gene itself. Without that information, no enabler mutations could be identified, so it was also classified as a putative *de novo* gene and its status cannot be confirmed.

**Figure 4.**
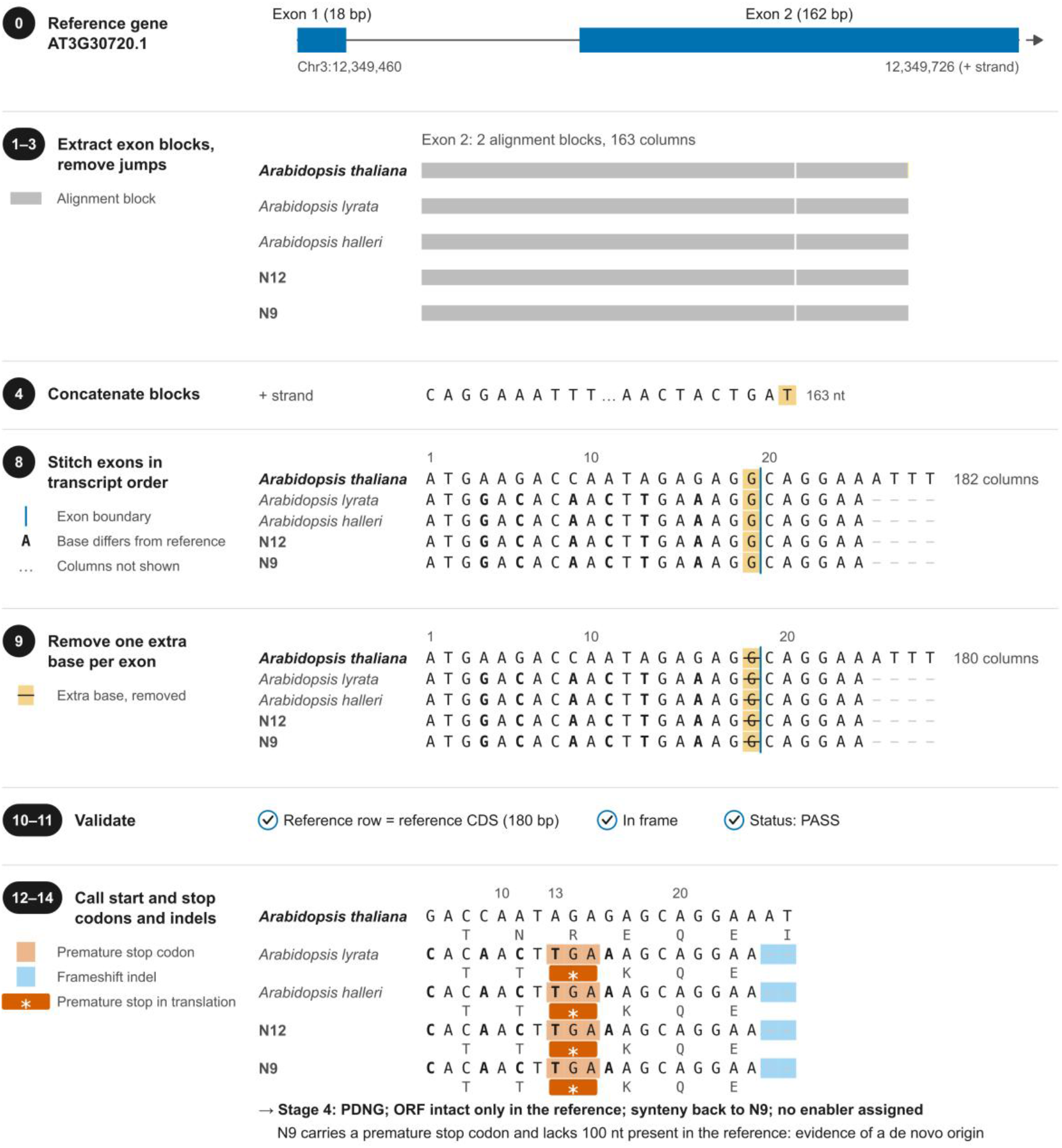
*A. thaliana QQS* gene (*AT3G30720*.*1*) from the Brassicaceae run. Steps and symbols are as in Fig. 3. The gene contains two exons on the plus strand of chromosome 3 (18 and 162 bp). No species were filtered out in steps 1–3. *A. lyrata, A. halleri* and the reconstructed ancestors N12 and N9 carry a TGA premature stop codon at positions 13–15 (AGA in *A. thaliana*) and have a string of gaps from positions 25 to 124, the beginning of which is shown. The ORF is intact only in *A. thaliana*, and synteny extends back only to N9, its immediate ancestor, so *QQS* is classified as a putative *de novo* gene with no enabler assigned. The premature stop codon and missing sequence in N9, N12, *A. lyrata* and *A. halleri*, absent from *A. thaliana*, are nonetheless evidence of *de novo* origination. For further explanation of each step see the Genomes2Genes Overview.

## Conclusions

Progressive Cactus (PC) has enabled whole-genome alignments of hundreds of large assemblies, ushering in a new era of large-scale comparative analyses. However, the development of new pipelines and tools to verify, parse and analyze PC whole-genome alignments remains essential to fully leverage the applications of this tool and its capacity to answer genomic and evolutionary questions. The Genomes2Genes pipeline is a new conduit that addresses the shortcomings of the currently available PC outputs and analysis tools. It has the potential to be applied to any question or analysis relating to comparative genomics, genes, and evolutionary biology. Different components of the pipeline can be used to conduct different studies, and outputs from across the pipeline can be incorporated into current and specialized investigations. Rigorous verification of the scripts and their outputs validated the computational accuracy of these analyses. The applications of PC and its whole-genome alignments continue to expand, and Genomes2Genes, presented here, extends them.

## Acknowledgments

We thank the members of the Casola laboratory for feedback on earlier versions of this manuscript.

## Statements and Declarations

### Author Contributions

S.C.S. designed and developed the pipeline and wrote the manuscript. C.C. designed and supervised the project.

### Ethics Approval

Not applicable.

### Consent to Participate

Not applicable.

### Data Availability

The BED files for the Brassicaceae run are in the bed_files directory of https://github.com/sierras64/Genomes2Genes-Brassicaceae. The Progressive Cactus whole-genome alignment is available at https://doi.org/10.5281/zenodo.22137464.

### Code Availability

Genomes2Genes is available at https://github.com/sierras64/Genomes2Genes-Brassicaceae

### Use of Large Language Models

ChatGPT (OpenAI) and Claude (Anthropic), through Claude Code, were used to assist with writing, running and debugging analysis and plotting scripts, checking computed values against the source data, searching the literature and editing the manuscript text for clarity and consistency. All code, analyses, figures and text were reviewed and verified by the authors, who take full responsibility for the content of this article.

## References

Andrews G, Fan K, Pratt HE et al (2023) Mammalian evolution of human cis-regulatory elements and transcription factor binding sites. Science 380:eabn7930. 10.1126/science.abn7930

Arendsee Z, Li J, Singh U et al (2019) fagin: synteny-based phylostratigraphy and finer classification of young genes. BMC Bioinformatics 20:440. 10.1186/s12859-019-3023-y

Armstrong J, Hickey G, Diekhans M et al (2020) Progressive Cactus is a multiple-genome aligner for the thousand-genome era. Nature 587:246–251. 10.1038/s41586-020-2871-y

Casola C (2018) From de novo to ‘de nono’: the majority of novel protein coding genes identified with phylostratigraphy are old genes or recent duplicates. Genome Biol Evol 10:2906–2918. 10.1093/gbe/evy231

Casola C, Luria V, Vakirlis N, Zhao L (2025) De Novo Genes: Current Status and Future Goals. Genome Biol Evol 17:evaf230. 10.1093/gbe/evaf230

Casper J, Speir ML, Raney BJ et al (2026) The UCSC Genome Browser database: 2026 update. Nucleic Acids Res 54:D1331–D1335. 10.1093/nar/gkaf1250

Christmas MJ, Kaplow IM, Genereux DP et al (2023) Evolutionary constraint and innovation across hundreds of placental mammals. Science 380:eabn3943. 10.1126/science.abn3943

Claros MG, Bautista R, Guerrero-Fernández D et al (2012) Why assembling plant genome sequences is so challenging. Biology (Basel) 1:439–459. 10.3390/biology1020439

Cock PJA, Antao T, Chang JT et al (2009) Biopython: freely available Python tools for computational molecular biology and bioinformatics. Bioinformatics 25:1422–1423. 10.1093/bioinformatics/btp163

Earl D, Nguyen N, Hickey G et al (2014) Alignathon: a competitive assessment of whole-genome alignment methods. Genome Res 24:2077–2089. 10.1101/gr.174920.114

Feng S, Stiller J, Deng Y et al (2020) Dense sampling of bird diversity increases power of comparative genomics. Nature 587:252–257. 10.1038/s41586-020-2873-9

Fiddes IT, Armstrong J, Diekhans M et al (2018) Comparative Annotation Toolkit (CAT)— simultaneous clade and personal genome annotation. Genome Res 28:1029–1038. 10.1101/gr.233460.117

Hickey G, Paten B, Earl D, Zerbino D, Haussler D (2013) HAL: a hierarchical format for storing and analyzing multiple genome alignments. Bioinformatics 29:1341–1342. 10.1093/bioinformatics/btt128

Hickey G, Monlong J, Ebler J et al (2024) Pangenome graph construction from genome alignments with Minigraph-Cactus. Nat Biotechnol 42:663–673. 10.1038/s41587-023-01793-w

Jin GT, Xu YC, Hou XH et al (2025) A de novo Gene Promotes Seed Germination Under Drought Stress in Arabidopsis. Mol Biol Evol 42:msae262. 10.1093/molbev/msae262

Knowles DG, McLysaght A (2009) Recent de novo origin of human protein-coding genes. Genome Res 19:1752–1759. 10.1101/gr.095026.109

Krasheninnikova K, Diekhans M, Armstrong J et al (2020) halSynteny: a fast, easy-to-use conserved synteny block construction method for multiple whole-genome alignments. GigaScience 9:giaa047. 10.1093/gigascience/giaa047

Lewin HA, Robinson GE, Kress WJ et al (2018) Earth BioGenome Project: sequencing life for the future of life. Proc Natl Acad Sci USA 115:4325–4333. 10.1073/pnas.1720115115

Li L, Foster CM, Gan Q et al (2009) Identification of the novel protein QQS as a component of the starch metabolic network in Arabidopsis leaves. Plant J 58:485–498. 10.1111/j.1365-313X.2009.03793.x

Li L, Wurtele ES (2015) The QQS orphan gene of Arabidopsis modulates carbon and nitrogen allocation in soybean. Plant Biotechnol J 13:177–187. 10.1111/pbi.12238

Liao WW, Asri M, Ebler J et al (2023) A draft human pangenome reference. Nature 617:312–324. 10.1038/s41586-023-05896-x

McKinney W (2010) Data structures for statistical computing in Python. In: van der Walt S, Millman J (eds) Proceedings of the 9th Python in Science Conference. pp 56–61. 10.25080/Majora-92bf1922-00a

Paten B, Earl D, Nguyen N et al (2011) Cactus: Algorithms for genome multiple sequence alignment. Genome Res 21:1512–1528. 10.1101/gr.123356.111

Peng J, Zhao L (2024) The origin and structural evolution of de novo genes in Drosophila. Nat Commun 15:810. 10.1038/s41467-024-45028-1

Rhie A, McCarthy SA, Fedrigo O et al (2021) Towards complete and error-free genome assemblies of all vertebrate species. Nature 592:737–746. 10.1038/s41586-021-03451-0

Silveira AB, Trontin C, Cortijo S et al (2013) Extensive natural epigenetic variation at a de novo originated gene. PLoS Genet 9:e1003437. 10.1371/journal.pgen.1003437

Song B, Buckler ES, Stitzer MC (2024) New whole-genome alignment tools are needed for apping into plant diversity. Trends Plant Sci 29:355–369. 10.1016/j.tplants.2023.08.013

Song Y, Yuan Z, Fang C et al (2026) A genomic compendium of hundreds of teleost fishes reveals their evolutionary landscape. Innovation (Camb) 7:101177. 10.1016/j.xinn.2025.101177

Sullivan PF, Meadows JRS, Gazal S et al (2023) Leveraging base-pair mammalian constraint to understand genetic variation and human disease. Science 380:eabn2937. 10.1126/science.abn2937

Tang D, Jia Y, Zhang J et al (2022) Genome evolution and diversity of wild and cultivated potatoes. Nature 606:535–541. 10.1038/s41586-022-04822-x

Tange O (2011) GNU Parallel: the command-line power tool. ;login: The USENIX Magazine 36:42–47

Vakirlis N, Carvunis AR, McLysaght A (2020) Synteny-based analyses indicate that sequence divergence is not the main source of orphan genes. eLife 9:e53500. 10.7554/eLife.53500

Van Oss SB, Carvunis AR (2019) De novo gene birth. PLoS Genet 15:e1008160. 10.1371/journal.pgen.1008160

Walden N, German DA, Wolf EM et al (2020) Nested whole-genome duplications coincide with diversification and high morphological disparity in Brassicaceae. Nat Commun 11:3795. 10.1038/s41467-020-17605-7

Weisman CM, Murray AW, Eddy SR (2020) Many, but not all, lineage-specific genes can be explained by homology detection failure. PLoS Biol 18:e3000862. 10.1371/journal.pbio.3000862

Wendel JF, Jackson SA, Meyers BC, Wing RA (2016) Evolution of plant genome architecture. Genome Biol 17:37. 10.1186/s13059-016-0908-1

Wu Y, Li D, Hu Y et al (2023) Phylogenomic discovery of deleterious mutations facilitates hybrid potato breeding. Cell 186:2313–2328.e15. 10.1016/j.cell.2023.04.008

Yoo D, Rhie A, Hebbar P et al (2025) Complete sequencing of ape genomes. Nature 641:401–418. 10.1038/s41586-025-08816-3

Zhang X, Kaplow IM, Wirthlin M et al (2020) HALPER facilitates the identification of regulatory element orthologs across species. Bioinformatics 36:4339–4340. 10.1093/bioinformatics/btaa493

Zhang L, Ren Y, Yang T et al (2019) Rapid evolution of protein diversity by de novo origination in Oryza. Nat Ecol Evol 3:679–690. 10.1038/s41559-019-0822-5

Zhang Z, Zhang P, Ding Y et al (2025) Ancient hybridization underlies tuberization and radiation of the potato lineage. Cell 188:5249–5265.e15. 10.1016/j.cell.2025.06.034

Zhao L, Saelao P, Jones CD, Begun DJ. (2014) Origin and spread of de novo genes in Drosophila melanogaster populations. Science 343:769–72. 10.1126/science.1248286

Zoonomia Consortium (2020) A comparative genomics multitool for scientific discovery and conservation. Nature 587:240–245. 10.1038/s41586-020-2876-6

